# *Keep Garfagnina alive*. An integrated study on patterns of homozygosity, genomic inbreeding, admixture and breed traceability of the Italian Garfagnina goat breed

**DOI:** 10.1101/2020.04.16.044644

**Authors:** Christos Dadousis, Francesca Cecchi, Michela Ablondi, Maria Chiara Fabbri, Alessandra Stella, Riccardo Bozzi

## Abstract

The objective of this study was to investigate the genomic background of the Garfagnina (GRF) goat breed that faces the risk of extinction. In total, 48 goats genotyped with the Illumina CaprineSNP50 BeadChip were analyzed together with 214 goats belonging to 9 Italian breeds (~25 goats/breed) from the AdaptMap project [Argentata (ARG), Bionda dell’Adamello (BIO), Ciociara Grigia (CCG), Di Teramo (DIT), Garganica (GAR), Girgentana (GGT), Orobica (ORO), Valdostana (VAL) and Valpassiria (VSS)]. We estimated i) runs of homozygosity (ROH), ii) admixture ancestries and iii) traceability success via discriminant analysis on principal components (DAPC) based on cross-validation. For GRF, an excess of frequent ROH (more than 45% in the GRF samples analyzed) was detected on CHR 12 at, roughly 50.25-50.94Mbp (ARS1 assembly), spanned between the CENPJ (centromere protein) and IL17D (interleukin 17D) genes. The same area was also present in DIT, while the broader region (~49.25-51.94Mbp) was shared among the ARG, CCG, and GGT. Admixture analysis depicted the uniqueness of the GRF breed, with a small part of common ancestry shared with BIO, VSS, ARG and CCG breeds. The DAPC model resulted in a 100% assignment success. We hope this work will contribute to the efforts of preventing the GRF from extinction and to add value to all the socio-agro-economic factors related with the farming of the GRF breed.

## Introduction

Local breeds, that usually consist of a small number of animals, are increasingly recognized by E.U. action plans as a rule of rural land protection. There are several reasons for this. To name some, local breeds are i) rustic and resistant to their local environment, ii) they represent a significant economic resource and have been used for the manufacture of niche products, especially in mountainous regions, iii) they represent an important and usually unique gene bank that could be essential to address the future climate changes, or potential disease outbreaks [1], and iv) they play an important role for the preservation of the human cultural inheritance. Especially for small ruminants, that can adapt in marginal and difficult areas, another reason of major importance is the provision of eco-system services. In mountainous regions of the Mediterranean basin, grazing can be used as a measure of protection against avalanches in winter and fire outbreaks during the summer period. Grazing, apart from cost-effective is also a nonpolluting, nontoxic, nearly carbon-neutral and an effective technique against fire propagation. In this context, goat grazing has been proposed as an alternative and eco-friendly solution [2]. Moreover, climate change has been identified as an additional pressure to the sustainability of livestock systems (e.g., health and productivity) and local breeds provide with an alternative through the adaptiveness in the regions they are reared. Despite this, usually the low productivity of local unimproved breeds, and thereby low farmer’s income, endangers their existence.

Regarding goats (*Capra hircus*), their widespread presence and adaptation in a variety of agro-ecological conditions worldwide is well documented [3]. Goats, are closely related to the human kind history, since, together with sheep, cattle and pigs were from the earliest domesticated ungulates [3,4]. Nevertheless, based on the Domestic Animal Diversity Information System (DAD-IS) data, 21 goat breeds are extinct (18 from which were reared in the regions of Europe and Caucasus) and 41 are at critical situation (41 of which from Europe and Caucasus) [6]. In Italy, 3 goat breeds are extinct and 12 are marked in a critical situation.

The Garfagnina breed (GRF) is one of those breeds that faces the risk of extinction. The latest recognition reported 1,468 animals spread in 29 different farms (ARAT, personal communication). The GRF is reared, mostly for dairy production, in central Italy, in the hills and mountains of the northwestern Tuscan Apennine area. The origins of this population are not clear. However, it is very likely that the breed was a result of crossings between native goats from Alps and from the Tuscan-Emilian Apennines. Moreover, the local breeders report that the population was reared for generations for its milk and meat production [4]. The breed is also closely linked to the production of typical products, such as the Controneria meat kid and the Caprino delle Apuane cheese. As it has been reported by Martini et al. [4], the milking of GRF goats is manual.

To support the management and conservation of the breed, and to provide support to the farmers and to the general region where the breed is reared (hills and mountains of the northwestern Tuscan Apennine area - central Italy), a few studies investigated various production characteristics [4,7], the adaptive profile (through physiological, haematological, biochemical and hormonal parameters) [8] and resistance to diseases [9,10] of GRF. Martini et al. [4] investigated various zootechnical characteristics of the GRF breed in comparison to other Italian and foreign goat breeds. Based on their results, the authors suggested the development of a breeding scheme based on pure bred animals. Nevertheless, no whole genome analysis has been conducted yet to investigate the genomic background of GRF and its ancestry. Genomic information, however, is essential for action measures to be taken for conservation purposes. The 50K SNP chip (http://www.goatgenome.org; [11]) released in 2013 together with the recent results of the AdaptMap project [3] offered this opportunity.

Hence, the objective of the present study was to investigate the genomic background of the Garfagnina breed, relative to the native Italian goat breeds included in the AdaptMap dataset. A unified procedure on admixture, runs of homozygosity, and discriminant analysis was applied with results depicting the unique genetic structure of the breed.

## Materials and methods

### Ethics statement

Garfagnina goats belonged to commercial farms and blood sampling was conducted by veterinarians. No invasive procedures were applied. Thus, in accordance to the 2010/63/EU guide and the adoption of the Law D.L. 04/03/2014, n.26 by the Italian Government, an ethical approval is not required in our study.

### Genomic data

Blood samples of forty eight female GRF goats were collected and animals were genotyped with the Illumina GoatSNP50 BeadChip (Illumina Inc., San Diego, CA) containing 53,347 Single Nucleotide Polymorpishms (SNPs) [12]. Genomic data of nine Italian autochthonous goat breeds, namely Argentata dell’Etna (ARG), Bionda dell’Adamello (BIO), Ciociara Grigia (CCG), Di Teramo (DIT), Garganica (GAR), Girgentana (GGT), Orobica (ORO), Valdostana (VAL) and Valpassiria (VSS) were downloaded from the online repository (https://datadryad.org/stash/dataset/doi:10.5061/dryad.v8g21pt) of the ADAPTmap project [3,13]. The breeds were selected based on the breed abbreviation on the plink fam file downloaded from the repository and the breed description (code and country) reported in Table 1 of [13]. The two datasets were merged and quality control was conducted in PLINK v1.9 [3, 4] on the final dataset based on the following criteria: 1) only autosomes were kept, ii) call rate per SNP >95% and ii) missing values per sample <10%. After editing, 260 samples and 48,716 SNP were retained (Table 1). The distribution of the SNP per chromosome (CHR) is presented in S1 Fig.

**Table 1.** Name of breeds, breed code and number of animals analyzed before (pre-QC) and after (post-QC) quality control per breed.

| Breed name | Breed code | No. pre-QC | No. post-QC |
| --- | --- | --- | --- |
| Argentata dell'Etna | ARG | 25 | 24 |
| Bionda dell'Adamello | BIO | 24 | 24 |
| Ciociara Grigia | CCG | 19 | 19 |
| Di Teramo | DIT | 24 | 24 |
| Garganica | GAR | 20 | 20 |
| Girgentana | GGT | 30 | 30 |
| Garfagnina | GRF | 48 | 48 |
| Orobica | ORO | 24 | 23 |
| Valdostana | VAL | 24 | 24 |
| Valpassiria | VSS | 24 | 24 |

### Runs of homozygosity

Analysis of runs of homozygosity (ROH) was conducted in the R (*v. 3.5.0*) package *detectRUNS v. 0.9.5* [16] using the consecutive method [17] that runs under the main function *consecutiveRUNS.run*. The required parameters were set to: i) minimum number of 15 SNPs/ROH, ii) 1 Mbp minimum length of ROH and iii) allow one heterozygous SNP within an ROH (to account for genotyping errors). In addition, ROH lengths were split into five classes (0-2, 2-4, 4-8, 8-16 and >16 Mbp). For each of the class and breed, descriptive statistics of ROH per breed, per chromosome, per SNP and per length class were estimated. Principal component analysis (PCA) was used to identify (dis)similarities among breeds, relative to the average number of ROH identified per chromosome. In addition, genomic inbreeding (F_ROH_) was calculated per breed. Regions with an excess of frequent ROH (≥ 45%) were detected and surrounding genes (1 Mbp up/downstream) were identified using the *Capra hircus* ARS1 (http://www.ensembl.org/index.html) and the variant effect predictor (https://www.ensembl.org/Tools/VEP) Ensembl databases.

### Population stratification and ancestry

PCA and admixture analysis were used to infer the presence of distinct populations based on the genomic data. The proportion of mixed ancestry in the breeds was assessed by the *ADMIXTURE 1.22* software [7, 8]. The number of ancestries (K) to be retained in the admixture analysis (K = 2 to 10) was evaluated via a 10-fold cross-validation (CV). The final selection on the number of ancestries was done by inspecting the CV error.

### Discriminant analysis of principal components

Discriminant analysis was applied to assess the traceability of the GRF goats using genomic data. To achieve this, the methodology of discriminant analysis of principal components (DAPC) [20] implemented in the R package *adegenet* [5, 10, 11] was adopted. In brief, DAPC is a 2-step approach: firstly, a PCA on the matrix of the genotypes is conducted and then, a small number of selected PCs (instead of the original SNP genotypes) is used as an input for the linear discriminant analysis (LDA). The selection on the optimal number of PCs to be further used in the LDA is done via cross-validation (CV) were the data is split in training and validation sets. For the selection of PCs the following criteria were implemented: i) 10-fold CV with 30 repetitions, ii) a maximum number of 300 PCs were tested, and iii) the number of PCs to be retained was based on number of PCs associated with the highest mean success. Three different scenarios of DAPC were applied as described below:

1. Scenario 1 (supervised learning). The full dataset was analysed simultaneously. In this scenario, all available data were used for model training and the discriminant functions were extracted based on all animals. This is not, however, a real case scenario, since the discriminant functions were developed utilizing the entire data set. The objective for a practical application is to identify an external individual membership to a group (i.e. external validation). Hence, two more scenarios were developed adopting a CV scheme also for the discriminant function.
2. Scenario 2 (semi-supervised learning). Assessment of correct assignment of GRF goats was done via a semi-supervised CV (CV_SS_). Five GRF goats were sampled representing the testing set of the DAPC analysis. The reference population was constituted by the rest of 43 GRF goats plus all the goats from the other breeds. The five GRF samples were classified in one of the 10 breeds presented in the reference population via the function *predict.dapc*. The procedure was repeated 10 times and results were averaged over the 10 repetitions.
3. Scenario 3 (unsupervised learning). Assignment of GRF goats in a breed but without the presence of any GRF goats in the reference population and model training (unsupervised CV; CV_US_). This scenario could also be viewed as a method to assess the genomic similarity of the GRF with the rest of the breeds (i.e., type of clustering). The approach was similar to Scenario 2 other than the testing population consisted of the entire GRF set and GRF samples had to be classified in one or more of the other 9 breeds. To increase the number of the tested samples in each round of the CV, 80% of the GRF breed was sampled. Moreover, to test for the effect of the size in training the model (TRN) in the assignment of the GRF, different proportions of the reference population were sampled (20, 30,…,90%) 10 times each, and results were averaged over the 10 repetitions. In other words, the size of the reference population varied between 42 to 91 goats. All nine breeds were present in each scenario and all GRF goats were used in this scenario.

It should be noted that the terms (semi/un)-supervised should not be confused with the terminology used in machine learning. The introduction of these terms has been used in the manuscript to distinguish among the three approaches that have been used in the DAPC analysis, and, although they are, up to a point, analogous with the same terms used in machine learning they are not identical.

## Results

### Runs of Homozygosity

Summary results of the detected ROH regions as total counts or averaged based on the number of samples per breed are presented in Table 2 and Fig 1, respectively. A relative high number of ROH was detected for GRF (n=2,450), with the highest number being for GGT (n=2,762) followed by ORO (=2,693), while the smallest was found for ARG (n=465). For GRF, the number of ROH per *Capra hircus* chromosome (CHI) varied from 35 (CHI25) to 158 (CHI1). The maximum length of ROH per chromosome was found on CHI1 (568,887,711bp) and the minimum on CHI23 (113,323,345bp). In general, the total length of ROH per CHR followed the same pattern of the total ROH number per CHR (Fig 2).

**Table 2.** Total number of runs of homozygosity (ROH) detected per breed.

| Breed | No. ROH |
| --- | --- |
| ARG | 465 |
| BIO | 814 |
| CCG | 496 |
| DIT | 813 |
| GAR | 730 |
| GGT | 2,762 |
| GRF | 2,450 |
| ORO | 2,693 |
| VAL | 1,613 |
| VSS | 768 |

**Fig 1.**
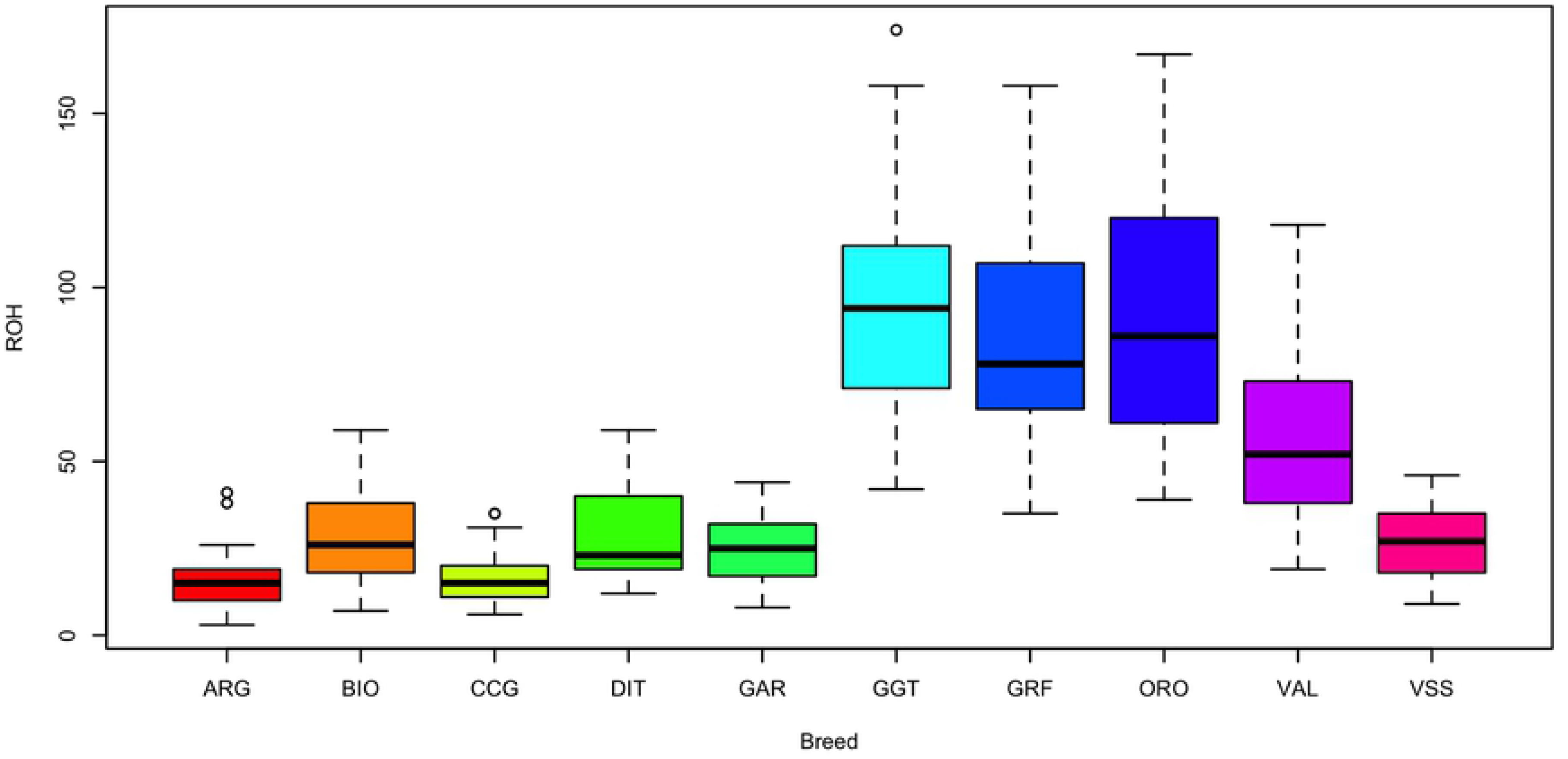
Average number of runs of homozygosity (ROH) detected per breed. ARG: Argentata dell’Etna; BIO: Bionda dell’Adamello; CCG: Ciociara Grigia; DIT: Di Teramo; GAR: Garganica; GGT: Girgentana; GRF: Garfagnina; ORO: Orobica; VAL: Valdostana and VSS: Valpassiria.

**Fig 2.**
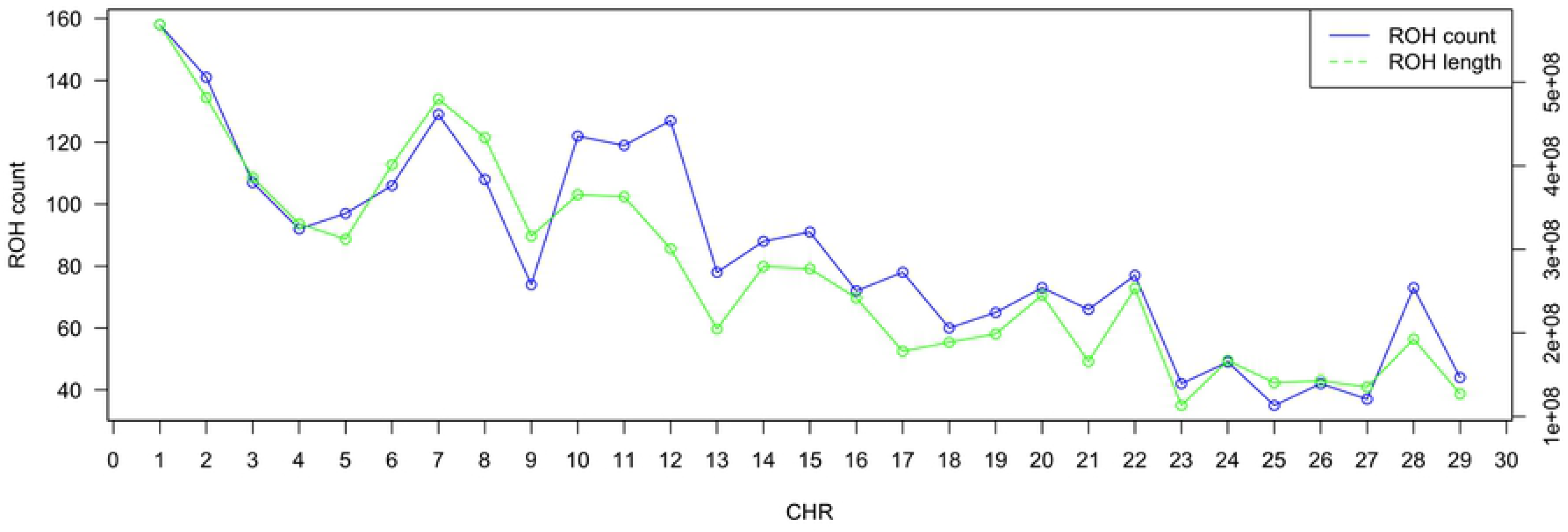
Number and length of runs of homozygosity (ROH) per chromosome (CHR) in Garfagnina breed.

For all breeds analyzed, except CCG and DIT, the number of ROH relative to the length on the genome was decreasing with an increased length (Fig 3α). In the DIT samples, the ROH were more frequent in length classes of 4-8 and 8-16 Mbp compared to the 2-4Mbp. The percentage of ROH with a length >16Mbp per breed varied between 0.89% to 13.53% for ORO and DIT, respectively. For small ROH length (<2Mbp) the proportion over the total number detected reached ~78% in ARG, while only 35% of ROH was observed for DIT. The pattern of ROH length class was similar among GRF, GGT and ORO with ~50% of the ROH having a length < 2Mbp, ~25% between 2-4Mbp, ~13% between 4-8Mbp, ~5% and ~2% between 8-16Mbp and > 16Mbp, respectively (S1 Table).

**Fig 3.**
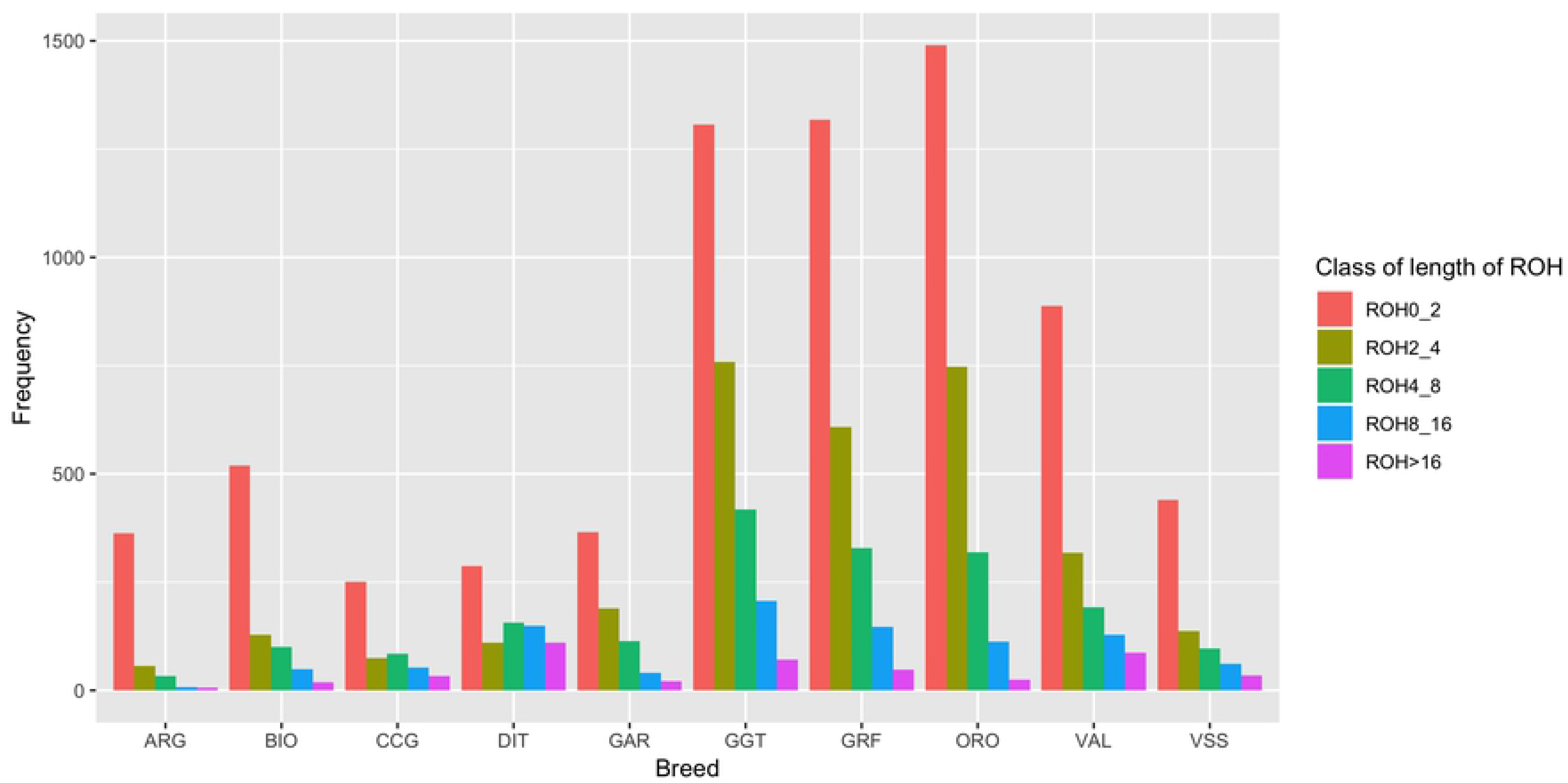
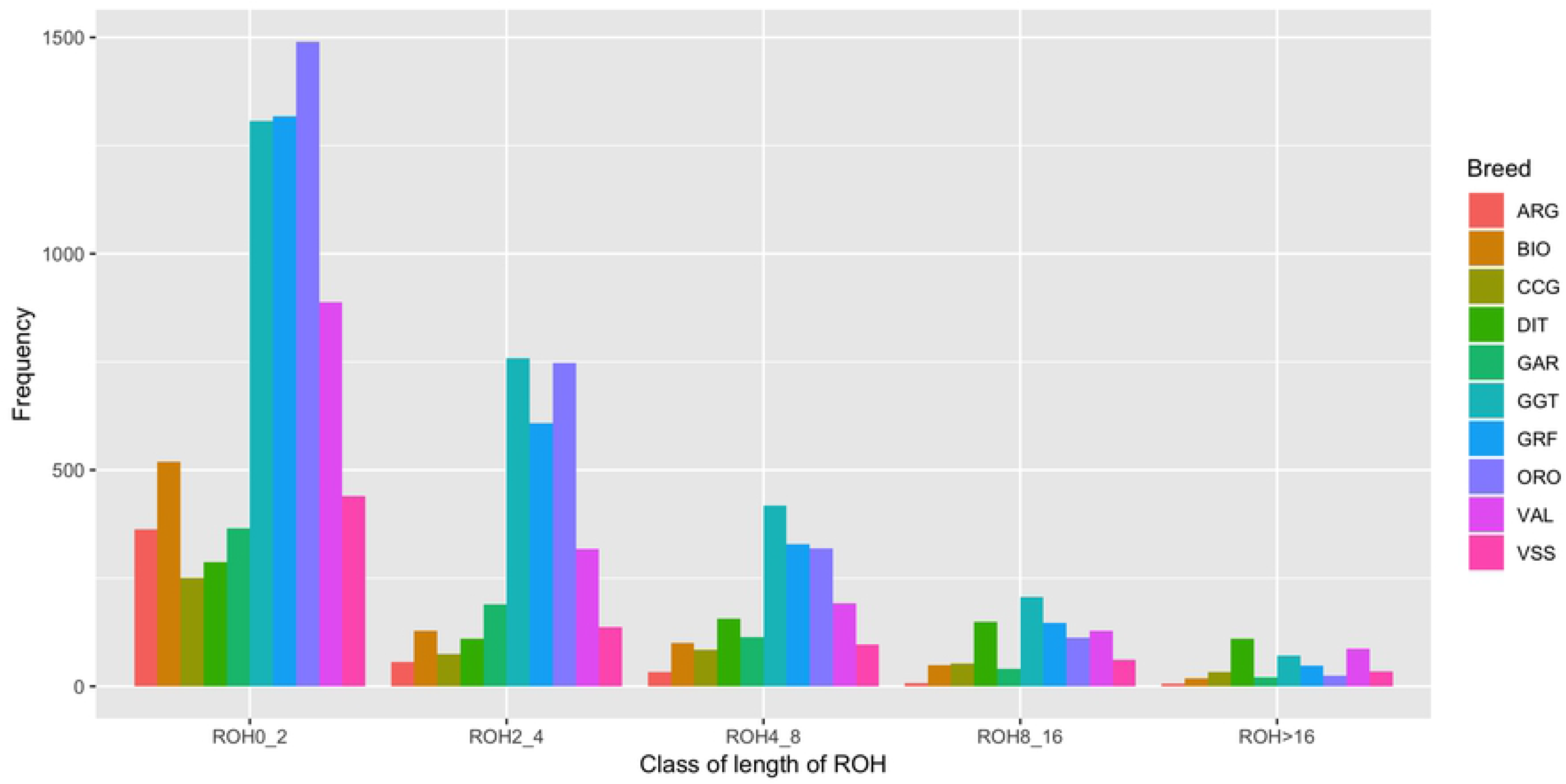
Frequency distribution of the number of runs of homozygosity (ROH) (a), in the breeds analyzed per length class, and (b) in different length classes per breed. ARG: Argentata dell’Etna; BIO: Bionda dell’Adamello; CCG: Ciociara Grigia; DIT: Di Teramo; GAR: Garganica; GGT: Girgentana; GRF: Garfagnina; ORO: Orobica; VAL: Valdostana and VSS: Valpassiria.

For GRF, an excess of frequent ROH (more than 45% in the GRF samples analyzed) was detected on CHI12, between ~34.6-35.3 Mbp (Fig 4, Table 3). In total, 14 SNP were contained in this region. The same excess area was also present in the DIT breed, while the broader region (~33,9-36.5 Mbp) was shared among the ARG, CCG, and GGT breeds. To identify similarities among breeds relative to the number of ROH per chromosome, a PCA was conducted on the average number of ROH identified per chromosome. In addition, a heatmap on the actual number of ROH per chromosome was produced. Both approaches placed GRF closer to ORO and GGT in respect to the rest of the breeds (Fig 5).

**Table 3.**
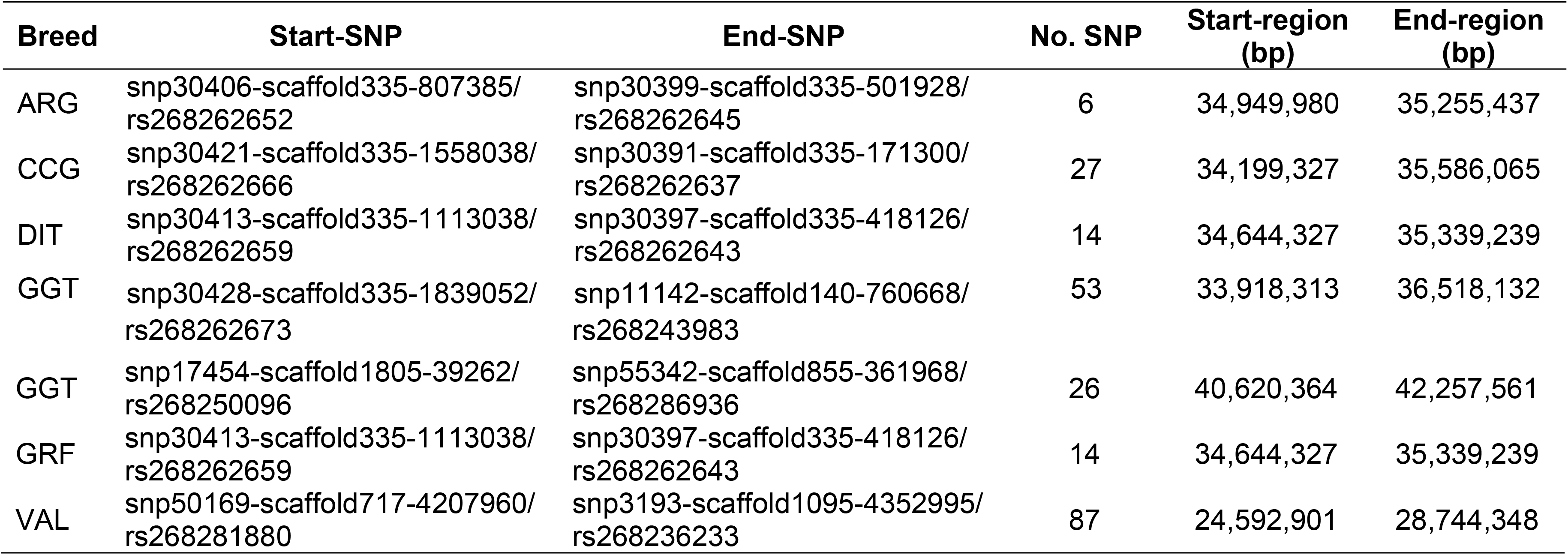
Most common (≥ 45% in each breed) runs of homozygosity (ROH) detected per breed on *Capra hircus* chromosome 12, with the start-end regions and number of SNP per ROH.

**Fig 4.**
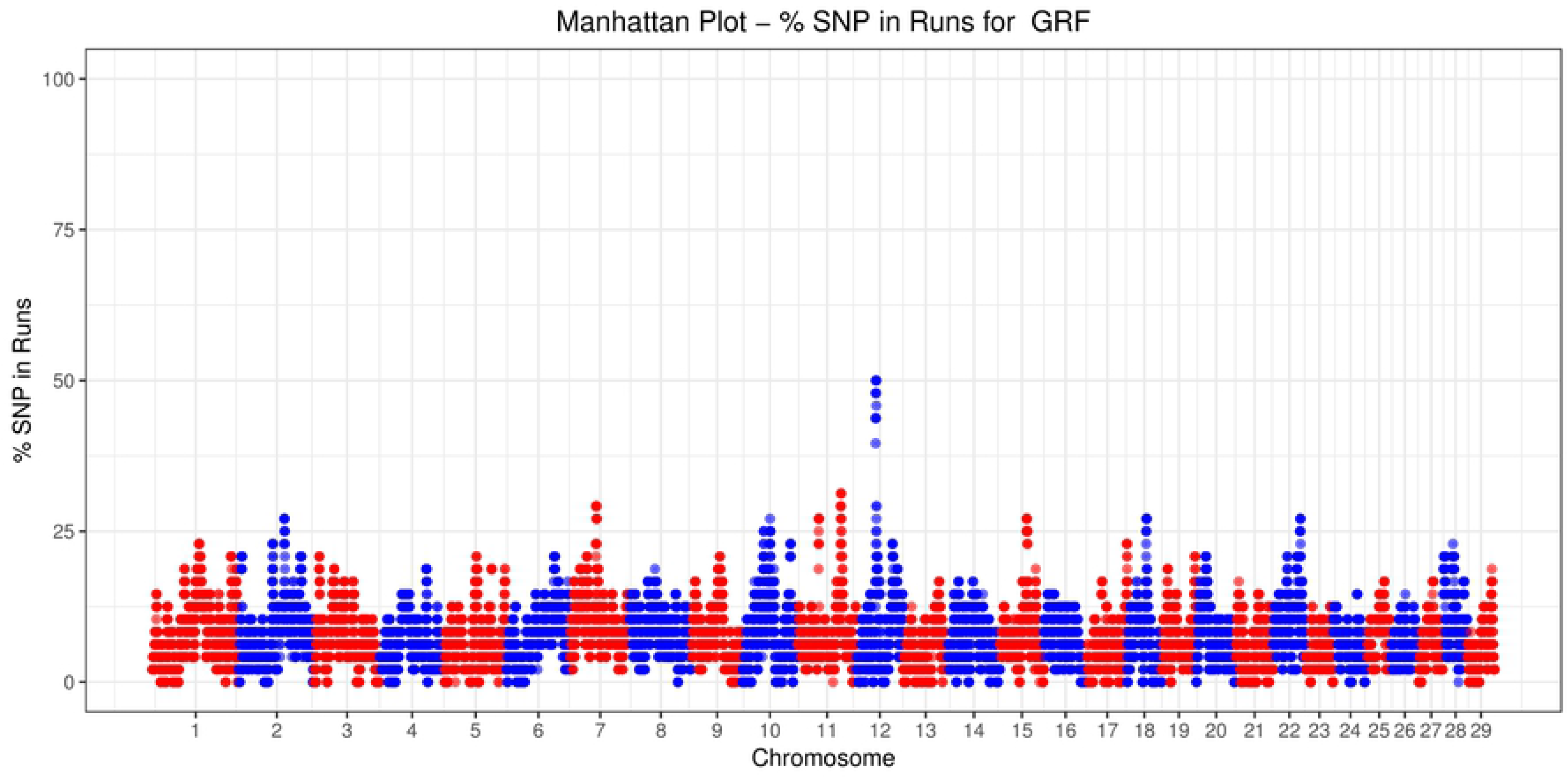
Number of times (%) each SNP was detected inside a run of homozygosity (ROH) in Garfagnina (GRF) goats.

**Fig 5.**
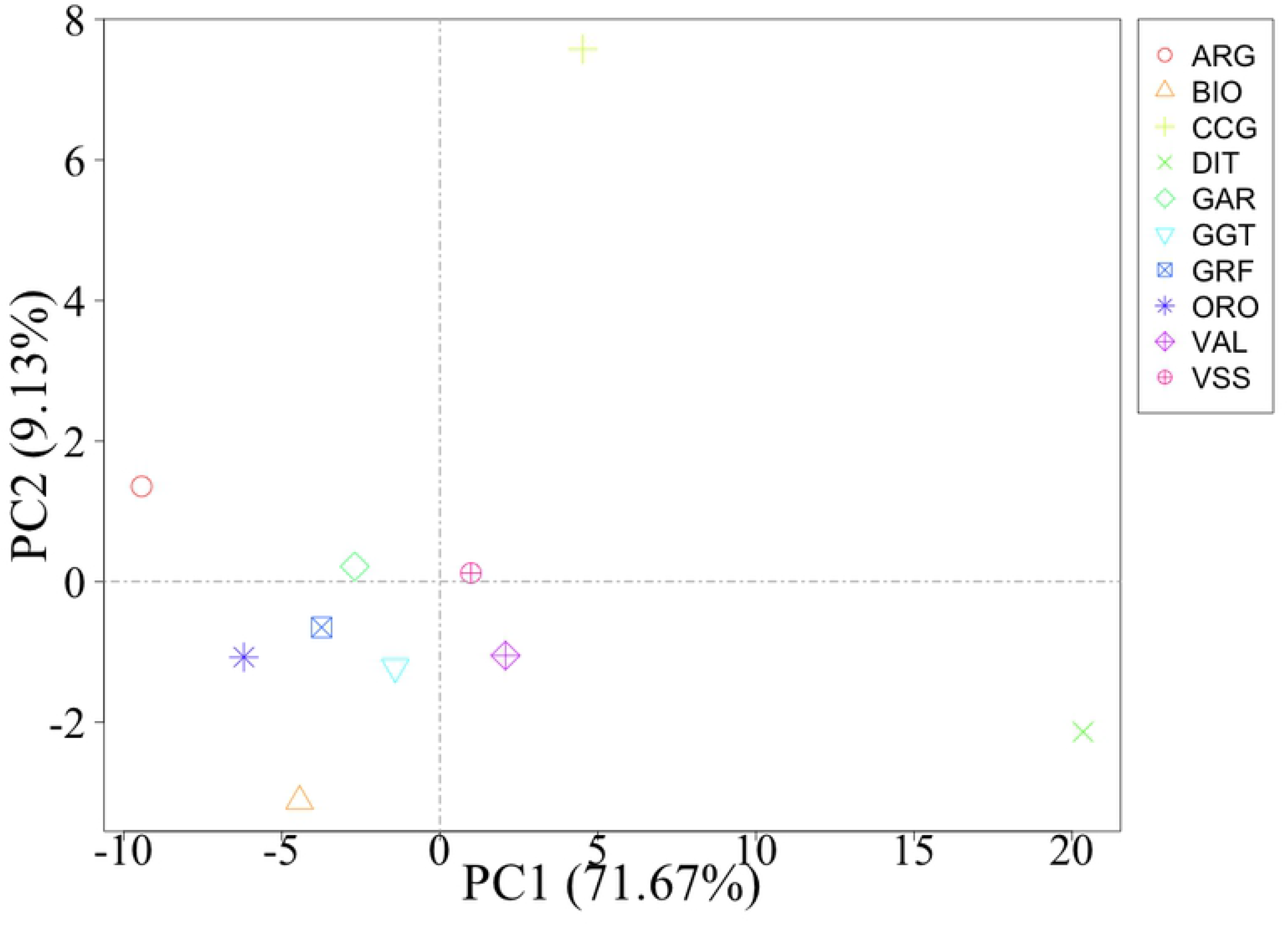
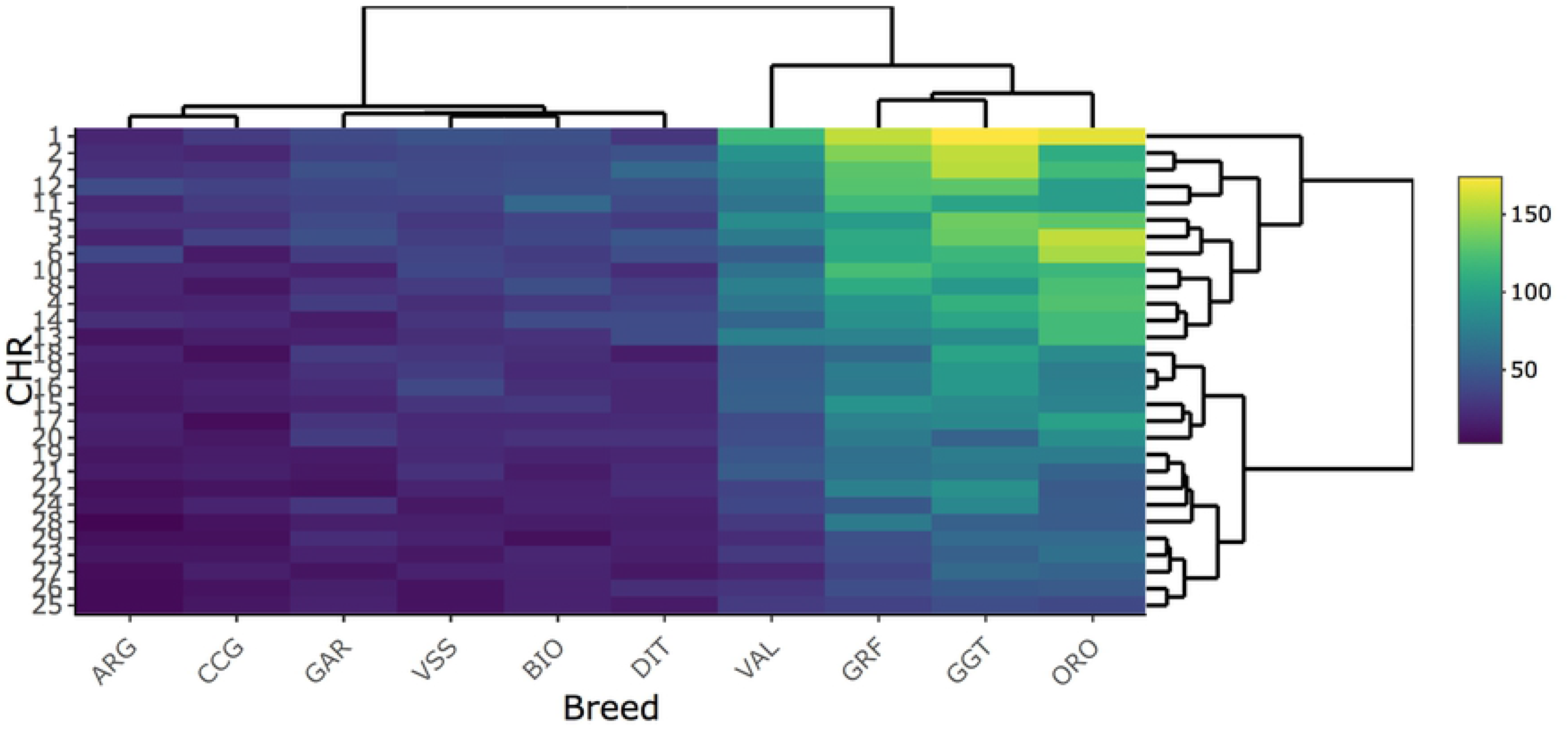
a) Scatterplot of principal component analysis conducted on the average runs of homozygosity identified per chromosome and breed; b) heatmap on the number of runs of homozygosity identified per chromosome and breed. ARG: Argentata dell’Etna; BIO: Bionda dell’Adamello; CCG: Ciociara Grigia; DIT: Di Teramo; GAR: Garganica; GGT: Girgentana; GRF: Garfagnina; ORO: Orobica; VAL: Valdostana and VSS: Valpassiria.

Genomic inbreeding coefficients (F_ROH_) were found intermediate for GRF compared to the rest of the breeds analyzed, with a mean value of 0.069 (Fig 6a, S2 Table). The highest values were observed for GGT (0.143) and ORO (0.137). Moreover, the distribution of F_ROH_ calculated per CHR was similar, with some high values (>0.5) observed for CHI7, 9, 16, 22 and 25 (Fig 6b).

**Fig 6.**
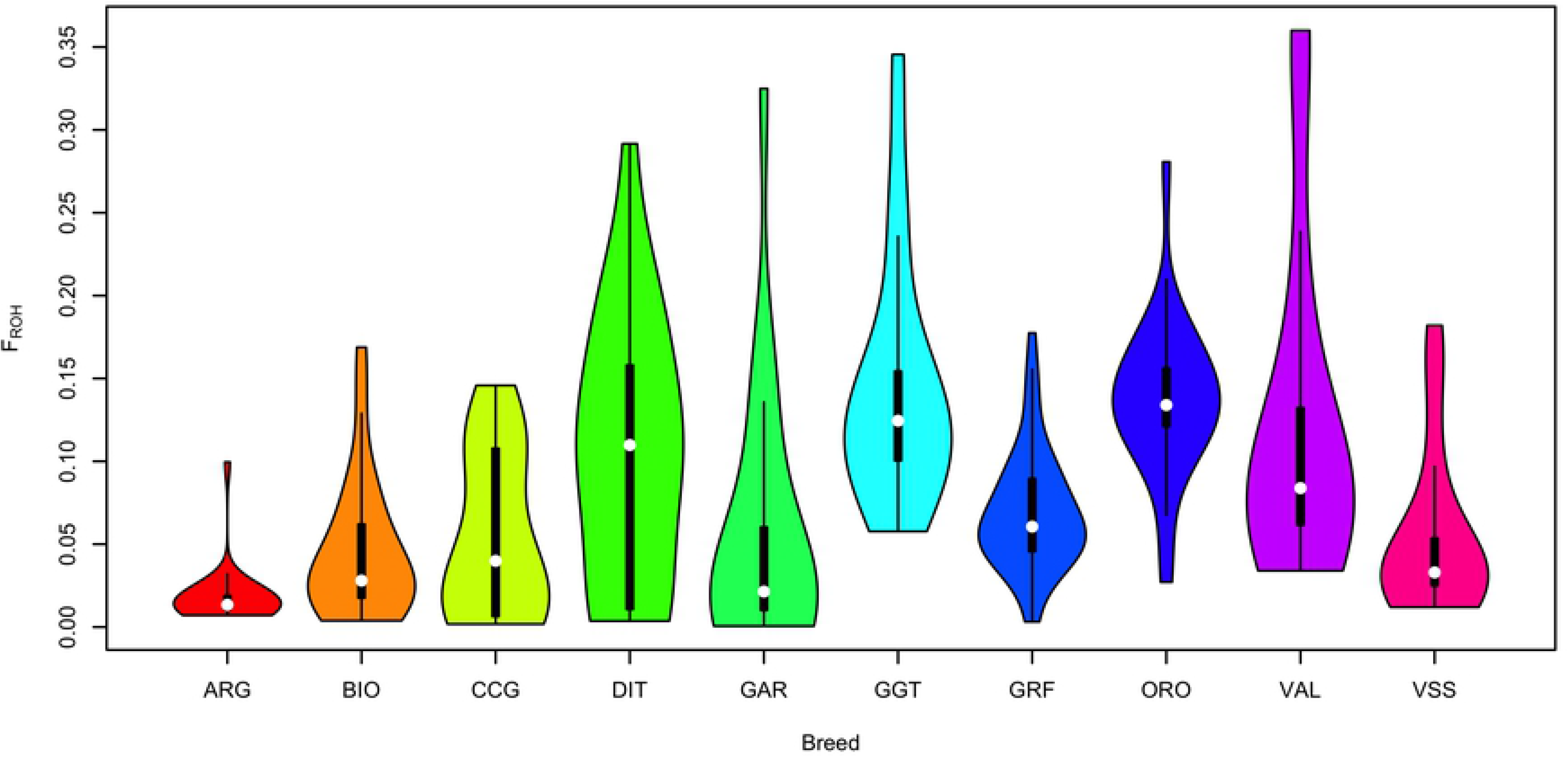
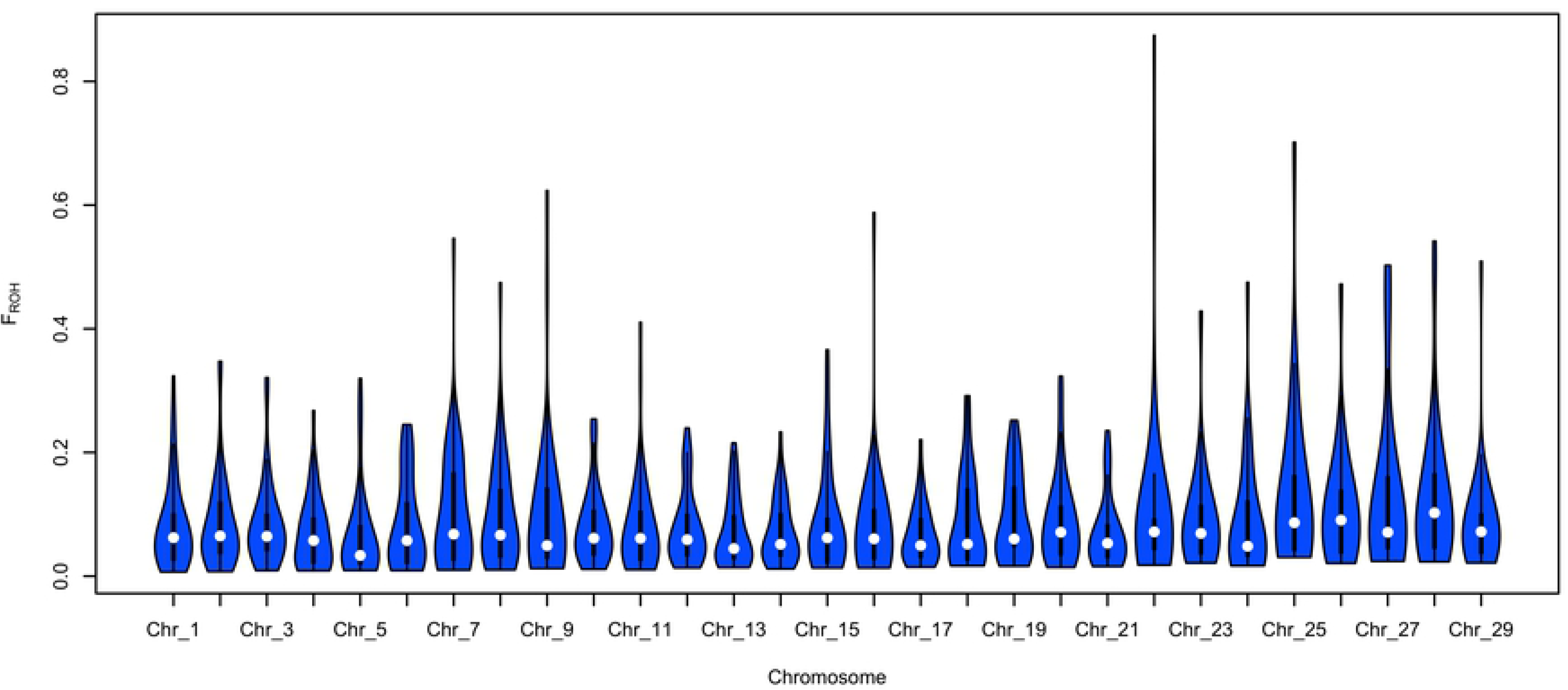
Summary of the genomic inbreeding coefficients (a) per breed and (b) of the Garfagnina breed per chromosome (Chr). ARG: Argentata dell’Etna; BIO: Bionda dell’Adamello; CCG: Ciociara Grigia; DIT: Di Teramo; GAR: Garganica; GGT: Girgentana; GRF: Garfagnina; ORO: Orobica; VAL: Valdostana and VSS: Valpassiria.

### Population Stratification and Ancestry

At a first step, a PCA was conducted on the complete data to visualize the general structure and relationships among breeds. The first axis distinguished GRF goats from ARG, CCG, DIT, GAR and GGT, while the second axis further separated GRF from the rest of the breeds (Fig 7a). An inspection of all the pairwise comparison between the first 10 axes (PCs) was carried out. By plotting the PC1 vs. PC6 (Fig 7b) four clusters were observed, namely: i) DIT, GAR, and GGT, ii) ARG and CCG, iii) BIO and VSS while iv) GRF was grouped together with ORO and VAL.

**Fig 7.**
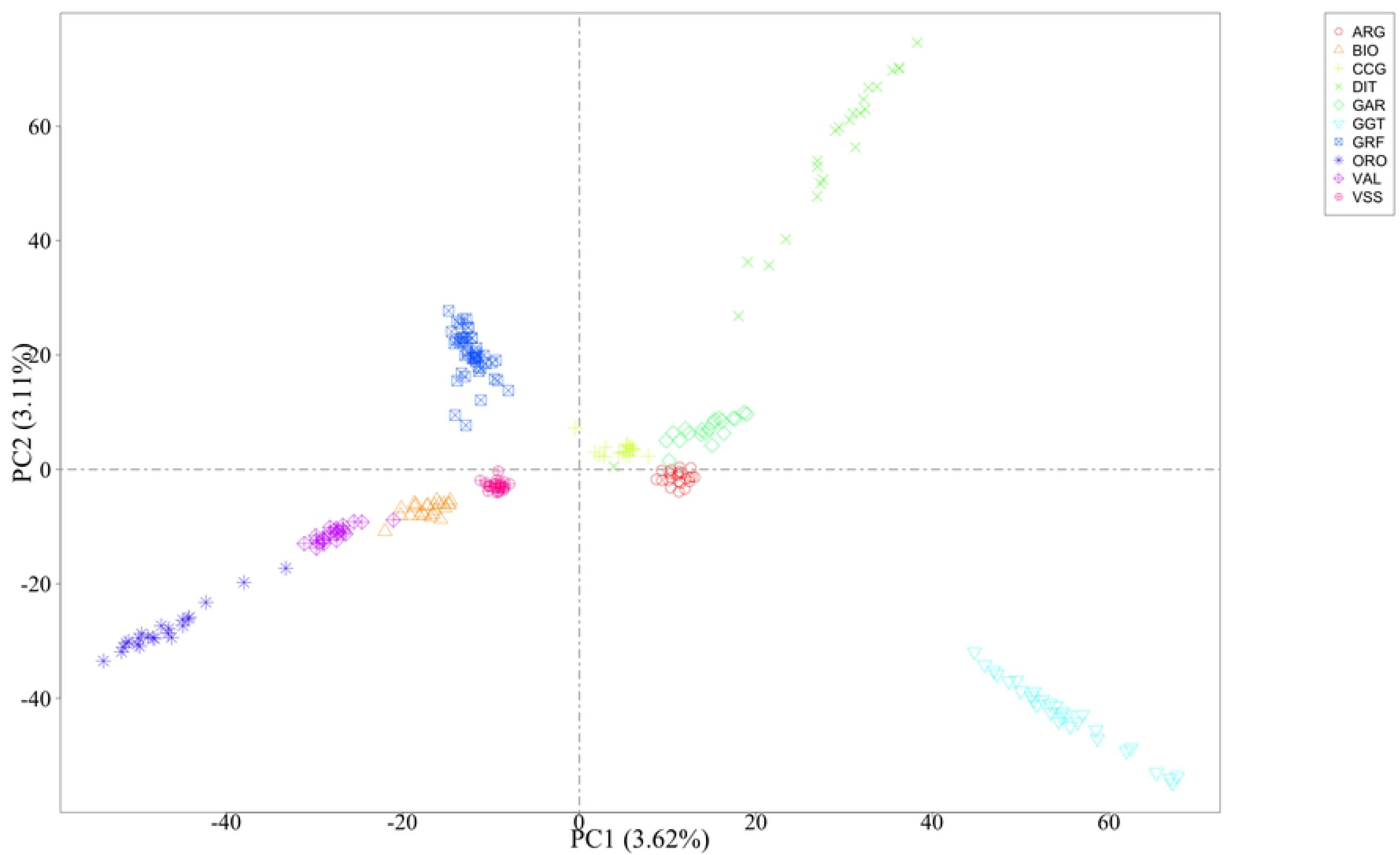
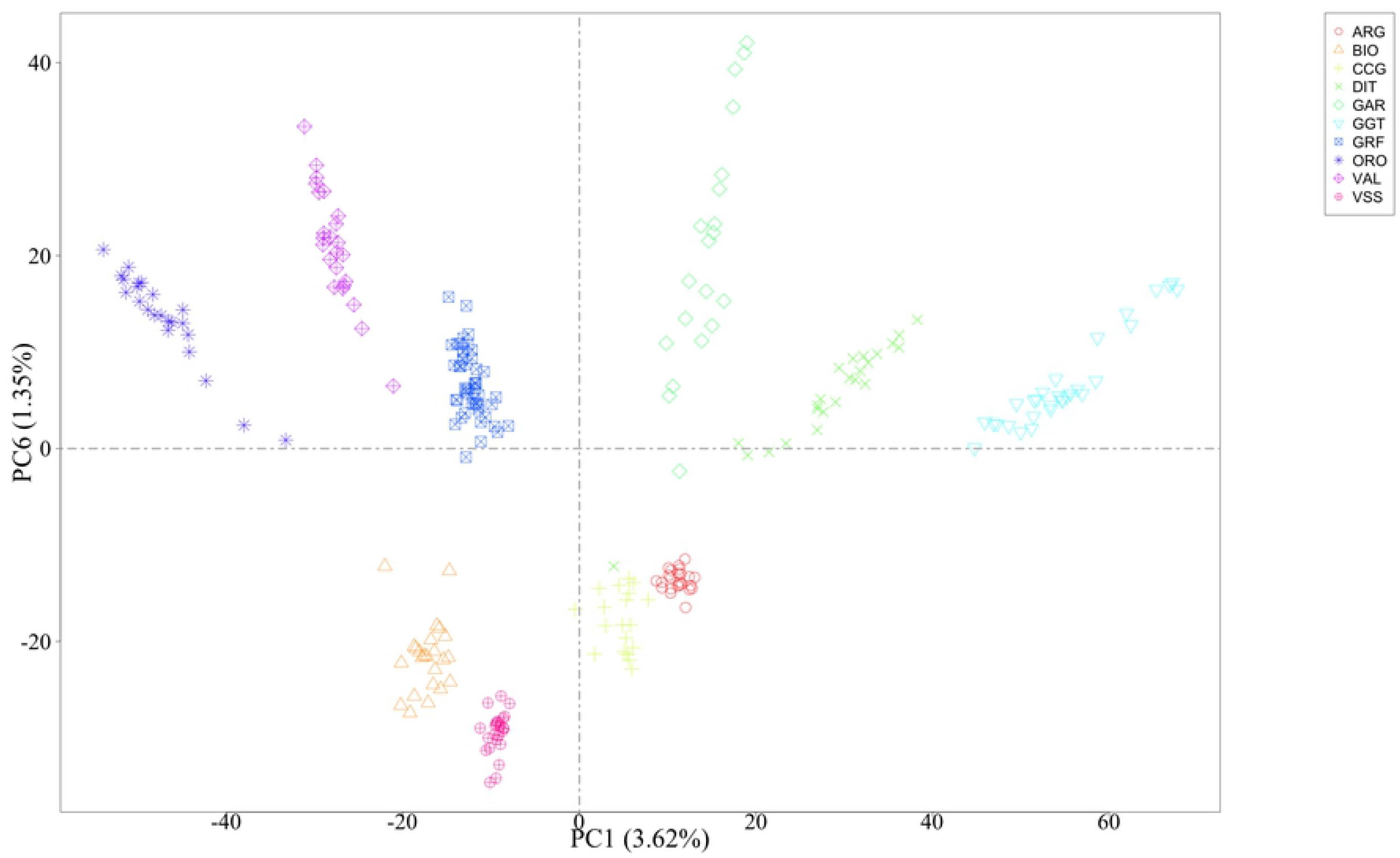
Scatterplot of (a) the first two and (b) first and sixth principal components. ARG: Argentata dell’Etna; BIO: Bionda dell’Adamello; CCG: Ciociara Grigia; DIT: Di Teramo; GAR: Garganica; GGT: Girgentana; GRF: Garfagnina; ORO: Orobica; VAL: Valdostana and VSS: Valpassiria.

An admixture analysis was conducted to complement with the PCA results. A varying number of group ancestries was investigated, from K=2 up to 10. The model with the minimum CV error was the one with eight group ancestries (S2 Fig). In general, the admixture results were in agreement with PCA, depicting the uniqueness of the GRF genome. At K=4, the GRF was shown almost as a breed-specific ancestry, sharing a small degree of ancestry primarily with ORO and VAL and further with GGT and DIT (Fig 8). At K=8, again, GRF had almost a breed-specific ancestry, with a small percentage of the GRF goats sharing co ancestry with i) BIO and VSS and ii) ARG and CCG and to a small extent with ORO, VAL, DIT and GGT. It should be noted that apart from GRF, group-specific ancestries, at least to a great extent, existed almost for all breeds but ARG, CCG, GAR and VSS.

**Fig 8.**
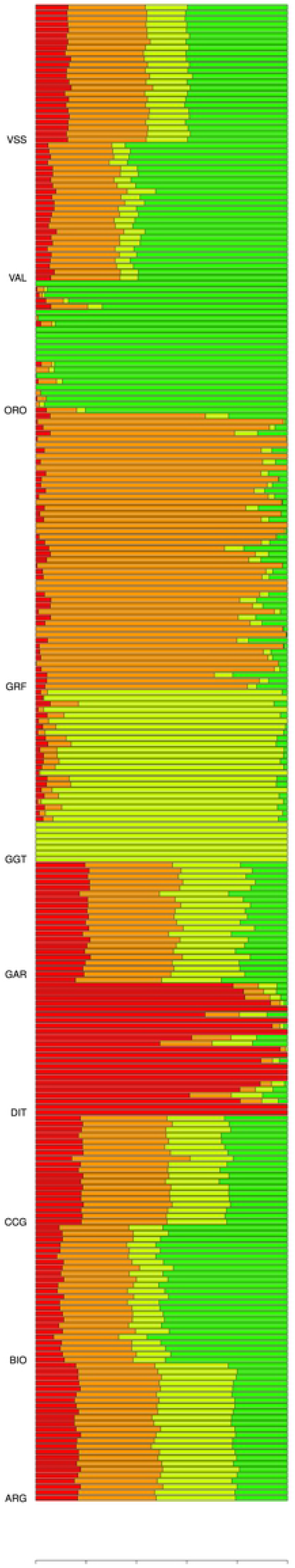
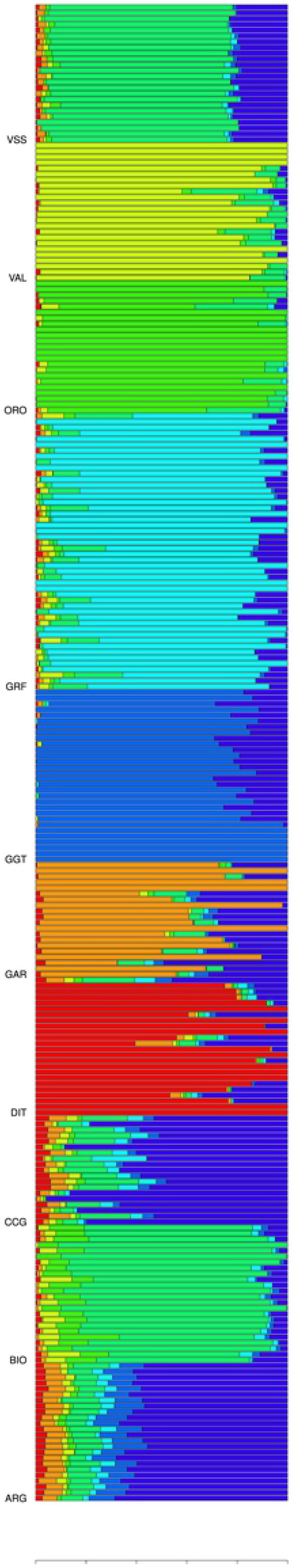
Admixture analysis with a) K=4 and b) K=8 coancestry groups. ARG: Argentata dell’Etna; BIO: Bionda dell’Adamello; CCG: Ciociara Grigia; DIT: Di Teramo; GAR: Garganica; GGT: Girgentana; GRF: Garfagnina; ORO: Orobica; VAL: Valdostana and VSS: Valpassiria.

### Discriminant Analysis of Principal Components

In the first scenario of DAPC, all data were used. The first 40 PCs, explaining ~35.75% of the total variability in the SNP data (S3 Fig), were used in the final DAPC model, resulting in an assignment success rate of 100% for all the GRF goats to its breed of origin. The pattern of the genetic diversity based on the DAPC is presented in Fig 9, where a clear genetic distance of GRF from the rest of the breeds can be observed.

**Fig 9.**
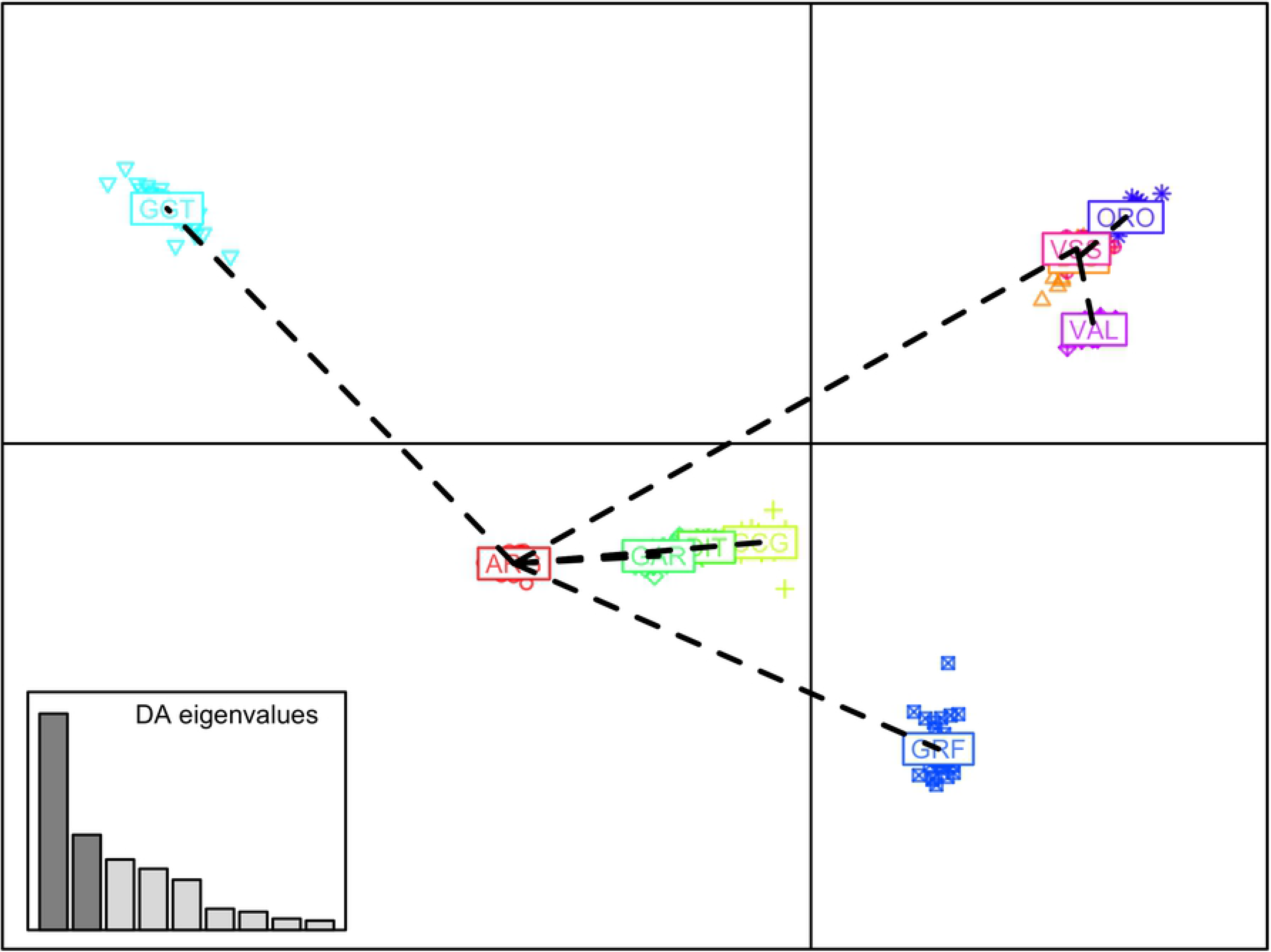
Scatterplot of the first two discriminant components of the DAPC. Breeds are presented by different colors and symbols. ARG: Argentata dell’Etna; BIO: Bionda dell’Adamello; CCG: Ciociara Grigia; DIT: Di Teramo; GAR: Garganica; GGT: Girgentana; GRF: Garfagnina; ORO: Orobica; VAL: Valdostana and VSS: Valpassiria.

An external validation scenario, that better reflects a practical application of the discriminant model, was further assessed. In the first analysis (CV_SS_ scenario), the GRF breed had representative animals in the reference population where the model of DAPC was developed. Also, in this case, a 100% correct classification of the GRF goats was observed (S3 Table). Interestingly, the classification of the GRF was invariant to the number of PCs selected (ranged between 10 to 70) in DAPC (S4 Table). In the second scenario (CV_US_) there were no representative GRF samples in training the model of DAPC. In that case the majority of animals were classified as CCG while few were assigned to DIT in some of the CV replicates (S5 Table). Similar results were obtained with an increased size of the reference population. In the majority of the scenarios, the GRF goats were classified either as CCG or DIT, where there were few cases in which GRF goats were also assigned as VSS, GAR or BIO, but in none of the cases as ORO, GGT or VAL (Fig 10).

**Fig 10.**
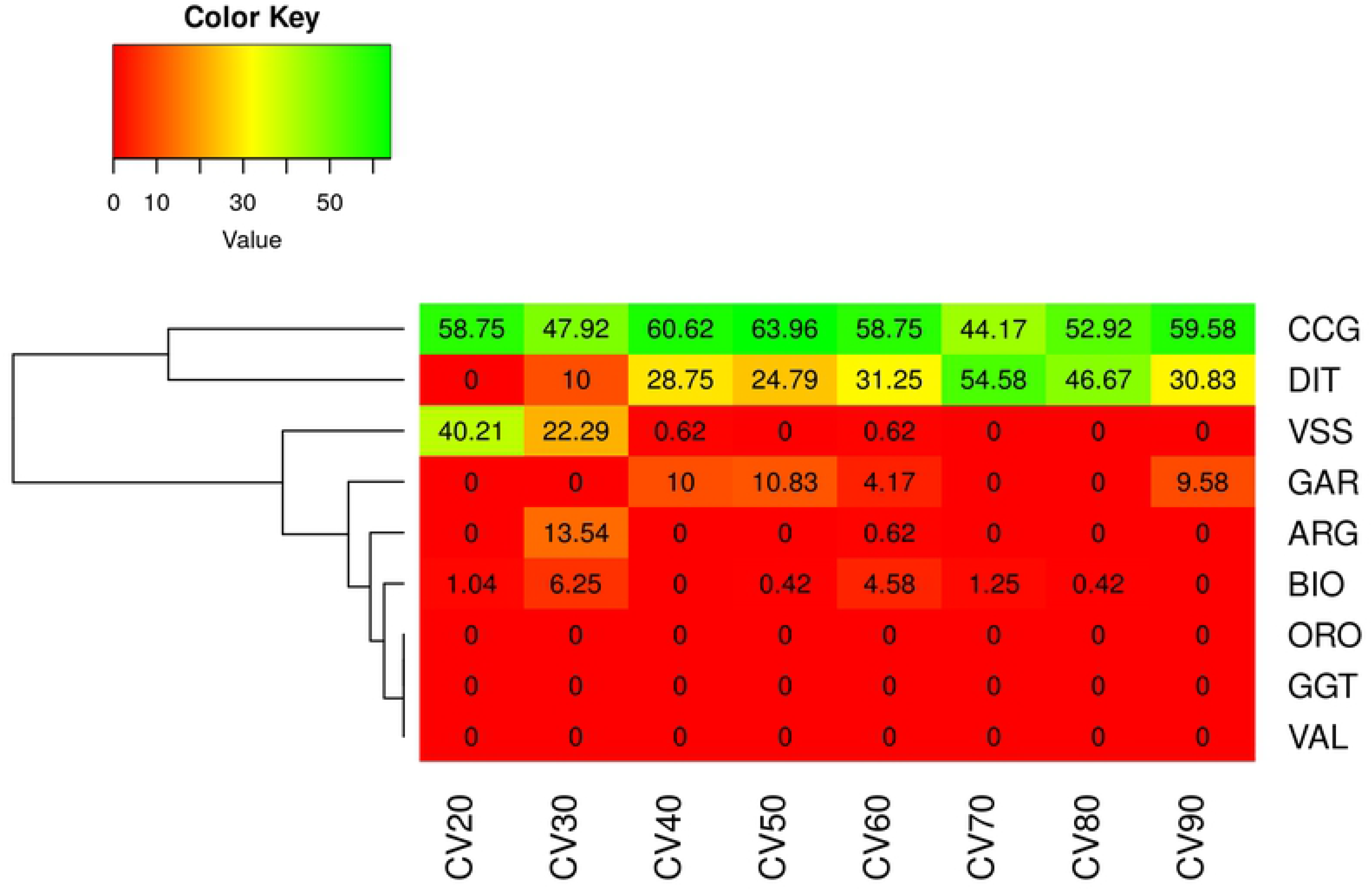
Percentage of assignment of the GRF goats in the CV_US_ scenario. Results were averaged over 10 replicates (CV) in each data subset (from 20 to 90%). CV_US_: unsupervised CV, where GRF breed had no representative goats in model training, hence GRF goats had to be classified in one of the rest 8 breeds; ARG: Argentata dell’Etna; BIO: Bionda dell’Adamello; CCG: Ciociara Grigia; DIT: Di Teramo; GAR: Garganica; GGT: Girgentana; GRF: Garfagnina; ORO: Orobica; VAL: Valdostana and VSS: Valpassiria.

## Discussion

The GRF breed is one of the Italian native goat breeds facing the risk of extinction, with a total number of registered animals lower than 1,500. Given the risk status of the breed, scientists have focused in a better description the GRF population. Characteristics on various zootechnical parameters of the breed, for instance, the milk total and fatty acid composition, milk coagulation properties and casein genotypes have been previously investigated [4], with authors encouraging the development of a purebred breeding scheme. Nevertheless, up to present, a whole-genome population analysis has not been carried out to study the GRF genome in terms of genetic diversity.

Following from the above, the present study aimed at describing the genomic profile of the GRF breed, relative to other Italian breeds for which genomic data were available. To achieve this, a sample of 48 genotyped GRF goats was merged with 214 genotyped goats coming from 9 Italian native breeds, 7 of them considered as dairy breeds and only 2 as meat breeds (ARG and VSS). These last have recently been analysed and presented from the AdaptMap project [3], hence we focused in presenting the results for the GRF breed. Our analysis was split in three parts, namely: i) runs of homozygosity, ii) principal component and admixture ancestry and iii) discriminant analysis.

### Runs of homozygosity

Runs of homozygosity can provide with a useful source of information on the historical background and the breeding management of a population. For instance, large ROH segments could be a result of recent intense selective breeding or of a potential bottleneck effect. As it has been highlighted in the work of Bertolini et al. in goats [23], crossbred populations tend to have smaller total ROH length and number compared to purebred populations. Same pattern was observed comparing unselected vs. selected populations undergoing breeding programs. Nevertheless, as it has been pointed out by [22,23] the 50k chip could not be considered adequate for an accurate detection of the small ROH, resulting in underestimation of small ROH hits. Despite this, our analysis was based on purebred goats, and thereby a smaller bias is expected. Results of ROH and F_ROH_ for the 9 Italian breeds of the AdaptMap project were in agreement with the estimates already reported [23]. Hence, our discussion on ROH is focused on highlighting the results on GRF in comparison to the rest of the 9 breeds analyzed.

The general pattern of ROH (i.e. in terms of total – and by chromosome – number and length of ROH) for GRF was similar to GGT and ORO. Moreover, an excess of frequent ROH was found for GRF (more than 45% in the GRF samples analyzed) on CHR 12 at, roughly, between 34.6-35.3 Mbp (Table 3). The same region was also detected in the DIT breed, while the broader region (~33,9-36.5 Mbp) was shared among the ARG, CCG, and GGT. Further, a search of genes presented in the top ROH region identified for GRF (~50.25-50.94Mbp, updated on the ARS1 assembly) and 1Mbp up-downstream (~49.25-51.94Mbp, updated on the ARS1 assembly) was carried out. Interestingly, the region ~49-52Mbp has been previously reported in goats [22–26]. It is worth noting that within this region lay the genes of the general gap junction protein family *GJA3* (gap junction protein alpha 3; ~50.642-50.644Mbp), *GJB2* (gap junction protein beta 2; ~50.675-50.676Mbp), *GJB6* (gap junction protein beta 6; ~50.694-50.695Mbp). The *GJB2* and *GJB6* are associated with the nervous system, hearing functions and ectodermal [24,25]. Moreover, the *SAP18* (Sin3A associated protein 18; ~51.136-51.141Mbp) that is related to gonad development [28], is also mapped in this region.

The narrow region of the detected top ROH runs for GRF on CHI12 was spanned between the *CENPJ* (centromere protein J; 50.23-50.27Mb) and the *IL17D* (interleukin 17D; 50.91-50.93 Mb). More precisely, the snp30397-scaffold335-418126 was found to be an intron of the *CENPJ* gene, while the snp30413-scaffold335-1113038 was downstream the *IL17D*. There are a series of studies that have linked the *CENPJ* with primary microcephaly in humans and in mice [28–31]. Moreover, *CENPJ* has been found to regulate in mouse the neurogenesis and the cilia disassembly in the developing corex [33]. Also in mouse, disruption of the *CENPJ* can cause the Seckel Syndrome [34].

Apart from the *CENPJ* and *IL17D* genes, within this genomic area, 9 more genes can be found, namely *PARP4* (poly(ADP-ribose) polymerase family member 4; ~50.29-50.35Mbp), *MPHOSPH8* (M-phase phosphoprotein 8; ~50.36-50.41Mbp), *PSPC1* (paraspeckle component 1; ~50.44-50.48Mbp), *ZMYM2* (zinc finger MYM-type containing 2; ~50.56-50.63Mbp), as well as the *CRYL1* (crystallin lambda 1; ~50.77-50.83Mbp) and the *IFT88* (intraflagellar transport 88; ~ 50.84-50.91Mbp).

Downstream this region, in 1Mbp expansion, the genes *ATP12A* (ATPase H+/K+ transporting non-gastric alpha2 subunit; ~50.08-50.11Mbp) and *RNF17* (ring finger protein 17; ~50.11-50.23Mbp) are located. Moreover, upstream the region there are mapped also the *EEF1AKMT1* (EEF1A lysine methyltransferase 1; ~50.94-50.95Mbp), *LATS2* (large tumor suppressor kinase 2; ~51.07-51.09Mbp), *ZDHHC20* (zinc finger DHHC-type containing 20; ~51.16-51.22Mbp), *MICU2* (mitochondrial calcium uptake 2; ~51.23-51.28Mbp), and the *FGF9* (fibroblast growth factor 9; ~51.34-51.37Mbp).

### Population Stratification and Ancestry

Two approaches, complementary to each other, have been used to infer the GRF relationships with 9 native Italian breeds, namely principal component and admixture analysis. PCA is widely used to identify structure in the data and to distinguish between groups of the samples. In that sense, the objective of PCA is to summarize (dis)similarities over the different groups in the data rather than the individual itself. On the other hand, admixture is focusing on the individuals; it provides with probabilities for each individual to be clustered in one of the pre-defined group ancestries. As such, the two approaches could be viewed as complementary rather antagonistic to each other.

Both of the analysis confirmed the distinguished and unique genetic background of the GRF breed and results were in general agreement. More precisely, the PCA scatterplot of PC1 vs. PC2 (Fig 7a) placed the GRF closer to VSS, BIO and CCG. This finding was similar to the admixture analysis with K=8 (Fig 8b), which was the final model selected after CV comparison. At K=4 in admixture, GRF shared co-ancestry with the ORO and VAL breeds (Fig 8a). In PCA this relationship has been visualized by plotting PC1 vs. PC6 (Fig 7b).

As a further step, we investigated the potential of breed traceability based on genomic data. To this purpose, an LDA model was used, where genotypes were firstly transformed into PCs, and a small set of those (not more than 300) was fitted in LDA (DAPC analysis). The DAPC model was able to classify with 100% success the GRF goats to its breed of origin (S3 Table). Moreover, an unsupervised learning was applied, where the GRF had no representative samples in the reference population. Results were consistent with the PCA and admixture and assigned the majority of the GRF goats in the CCG breed with a small number of goats (varied between 4 to 6) assigned as DIT (S5 Table).

As mentioned above and in M&M, the primary step of the DAPC analysis is to select the number of PCs to be used in the discriminant model. Hence, a basic question was on how robust the DAPC could be considered relative to the number of PCs used. Our analysis showed that, although the assignment success was invariant to the number of PCs in the semi-supervised DAPC analysis (number of PCs varied between 10-70), a pattern was found in the case of the unsupervised model. More precisely, when the DAPC contained 40 PCs some of the goats were classified as DIT goats. In the rest of the cases, where 60 or 20 PCs were used, all of the GRF goats were assigned as CCG.

PCA analysis seems thus to individuate common ancestries between GRF goats and Alpine Arc goat breeds whereas the DAPC approach identifies similarities between GRF and the goat breeds from Central Italy. Both hypotheses are consistent with the history of the Tuscan goat populations that experienced migratory flows both from the North and from Central Italy. The genomic analysis confirms the hypothesis that the GRF breed is a result of crosses among goats from the Alpine Arc and Tuscan-Emilian Apennines regions. Nevertheless, at a great part, the GRF breed consists of a unique genetic pool, genetically distinguished from 9 other native Italian breeds, for which genomic information was available and analysed here. This, in turn, resulted in breed traceability with a 100% success rate after CV. To sum up, our analysis complements previous work on various zootechnical and adaptive characteristics [4,7,8] of the GRF population and provides with a more complete description of the breed.

## Conclusions

Our genomic analysis suggests a unique genetic pool of the Garfagnina breed, with small parts of common ancestry shared with Bionda dell’Adamello, Valpassiria, Argentata dell’Etna and Ciociara Grigia. Moreover, GRF can be successfully discriminated by the rest of the breeds analysed using genomic information with a success rate of 100%. This could help in breed traceability and controlling the amount of crossbreeding in the future. A ROH on CHI12 associated with the *CENPJ* gene should be further investigated in the population. We suggest conservation and breeding measures to be taken for the Garfagnina goat. We hope our work will add value to the GRF farming and the local region where the breed is reared.

## Supporting information

**S1 Fig. Number of SNPs per chromosome after quality control. The plot has been produced with the *synbreed R* package [15]**.

**S2 Fig. Cross-validation results for assessing the number of ancestry groups in the admixture analysis**.

**S3 Fig. Cross-validation results for the selection of principal components to be retained in the DAPC analysis**.

**S1 Table. Percentage of the number of runs of homozygosity per length class and breed**.

ARG: Argentata dell’Etna; BIO: Bionda dell’Adamello; CCG: Ciociara Grigia; DIT: Di Teramo; GAR: Garganica; GGT: Girgentana; GRF: Garfagnina; ORO: Orobica; VAL: Valdostana and VSS: Valpassiria.

**S2 Table. Descriptive statistics of the genomic inbreeding coefficients per breed**.

ARG: Argentata dell’Etna; BIO: Bionda dell’Adamello; CCG: Ciociara Grigia; DIT: Di Teramo; GAR: Garganica; GGT: Girgentana; GRF: Garfagnina; ORO: Orobica; VAL: Valdostana and VSS: Valpassiria.

**S3 Table. Assignment results of the Garfagnina goats in the semi-supervised cross-validation (CV_SS_) scenario with 10 repetitions**.

ARG: Argentata dell’Etna; BIO: Bionda dell’Adamello; CCG: Ciociara Grigia; DIT: Di Teramo; GAR: Garganica; GGT: Girgentana; GRF: Garfagnina; ORO: Orobica; VAL: Valdostana and VSS: Valpassiria.

**S4 Table. Number of principal components selected via cross-validation (CV) in the 1^st^ scenario of the DAPC analysis of the Garfganina breed (GRF)**.

CV_SS_: semi-supervised CV, where some GRF goats are present in model training; CV_US_: unsupervised CV, where GRF breed had no representative goats in model training.

**S5 Table. Assignment results of the Garfagnina goats in the external cross-validation (CV_US_) scenario with 10 repetitions**.

ARG: Argentata dell’Etna; BIO: Bionda dell’Adamello; CCG: Ciociara Grigia; DIT: Di Teramo; GAR: Garganica; GGT: Girgentana; GRF: Garfagnina; ORO: Orobica; VAL: Valdostana and VSS: Valpassiria.

